# Overcoming microenvironment-mediated chemoprotection through stromal galectin-3 inhibition in acute lymphoblastic leukemia

**DOI:** 10.1101/2021.09.24.461149

**Authors:** Somayeh S. Tarighat, Fei Fei, Eun Ji Joo, Hisham Abdel-Azim, Lu Yang, Huimin Geng, Khuchtumur Bum-Erdene, I. Darren Grice, Mark von Itzstein, Helen Blanchard, Nora Heisterkamp

## Abstract

Environmentally-mediated drug resistance in B-cell precursor acute lymphoblastic leukemia (BCP-ALL) significantly contributes to relapse. Stromal cells in the bone marrow environment protect leukemia cells by secretion of chemokines as cues for BCP-ALL migration towards, and adhesion to, stroma. Stromal cells and BCP-ALL cells communicate through stromal galectin-3. Here, we investigated the significance of stromal galectin-3 to BCP-ALL cells. We used CRISPR/Cas9 genome editing to ablate galectin-3 in stromal cells and found that galectin-3 is dispensable for steady-state BCP-ALL proliferation and viability. However, efficient leukemia migration and adhesion to stromal cells are significantly dependent on stromal galectin-3. Importantly, loss of stromal galectin-3 production sensitized BCP-ALL cells to conventional chemotherapy. We therefore tested novel carbohydrate-based small molecule compounds (Cpd14 and Cpd17) with high specificity for galectin-3. Consistent with results obtained using galectin-3-knockout stromal cells, treatment of stromal-BCP-ALL co-cultures inhibited BCP-ALL migration and adhesion. Moreover, these compounds induced anti-leukemic responses in BCP-ALL cells including a dose-dependent reduction of viability and proliferation, induction of apoptosis and, importantly, inhibition of drug resistance. Collectively, these findings indicate galectin-3 regulates BCP-ALL cell responses to chemotherapy through the interactions between leukemia cells and the stroma, and show that a combination of galectin-3 inhibition with conventional drugs can sensitize the leukemia cells to chemotherapy.

## Introduction

Relapse is a main cause of treatment failure for patients with B-cell precursor acute lymphoblastic leukemia (BCP-ALL)[1]. The tumor microenvironment is a major contributing factor to relapse because it regulates the migration, survival, proliferation and response to drug treatment in BCP-ALL cells [2, 3]. Notably, leukemia cell movement and adhesion to the right niche depends on the ability of these cells to migrate towards chemotactic cues such as SDF-1 produced by stromal cells [4]. Interfering with leukemia-microenvironment interactions has been proven effective in mobilizing leukemia cells away from their protective microenvironment, thus making them more accessible to standard chemotherapy [5-7].

We and others have modeled the interaction between BCP-ALL cells and stroma in an *ex vivo* tissue co-culture model with OP9 bone marrow stromal cells to identify interactions that promote leukemia cell survival. Mesenchymal bone marrow stromal cells synthesize and secrete particularly high amounts of galectin-3 (Gal3), a carbohydrate-binding protein with immunomodulatory activity (for example, [8-10]. We previously reported that Gal3 acts as a communicator between BCP-ALL cells and the stroma: it not only binds to the cell surface of BCP-ALL cells but is also actively internalized by them [11, 12]. Gal3 has a range of glycoconjugate ligands on the surface of cells, but intracellularly it also binds proteins [13] and regulates a variety of functions including growth and mRNA splicing [14-16].

Although we previously showed that Gal3 protein synthesized endogenously in BCP-ALL cells promotes their survival [11], the physiological effect of Gal3 produced by the tumor microenvironment on BCP-ALL cells was unknown. Here, we have approached this problem in two ways. Using Cas9/CRISPR we have knocked out Gal3 in bone marrow stromal cells to determine if any BCP-ALL functions are regulated by stromal-produced Gal3. We also used novel small molecule monosaccharide-based carbohydrate mimetics to examine the effect of drug-mediated Gal3 inhibition on BCP-ALL physiology. We conclude that Gal3 is a valid target for enhancing the effects of standard chemotherapy by interfering with the communication between BCP-ALL and stromal cells.

## Materials and methods

### Cells and cell culture

The murine OP9 bone marrow-derived stromal cell line (CRL-2749) was obtained from the American Type Culture Collection (Manassas, VA). The patient derived TXL2 (Ph-positive) and ICN3, ICN13, US7, ICN06, LAX39 and LAX40 (Ph-negative) pre-B acute lymphoblastic leukemias were described previously [17, 18]. BCP-ALL cells were grown on confluent irradiated or 10 μg/mL mitomycin-treated OP9 stromal cells using αMEM medium supplemented with 20% fetal bovine serum, 1% L-glutamine and 1% penicillin/streptomycin (Life Technologies, Grand Island, NY).

To isolate normal human BM MSC, screens used to filter products intended for bone marrow transplants and which are otherwise discarded were rinsed with 15 mL DOM media to dislodge cells. After removal of debris via a 40 μm cell strainer and centrifugation, cells were suspended in either αMEM + 20% FBS or in DOM medium [IMDM + 12.5% horse serum + 12.5% FBS]. 1% L-glu, 0.5% P/S and 0.1% β-mercaptoethanol were added and cells were cultured at normoxia on 10-cm dishes. Cells were passaged by trypsinization when confluent. MSC were defined as a population that is CD45^neg^ and positive for SSEA-4 [19] and CD271/LNGFR [20].

Primary BCP-ALL and normal bone marrow mononuclear cells were purified using Ficoll (#17144002, GE Healthcare, Pittsburgh, PA). Human specimen collection protocols were reviewed and approved by Children’s Hospital Los Angeles Institution Review Board (IRB) [Committee on Clinical Investigations] (CCI). Collections were in compliance with ethical practices and IRB approvals.

Synthesis and characterization of the taloside-based compounds 14 (Cpd 14; designated compound 1 in [21]) and 17 (Cpd 17; Bum-Erdene, manuscript submitted) are described in detail elsewhere. The compounds were dissolved in DMSO and stored at −80°C. Aliquots were not used more than three times. Nilotinib (AMN107) (Novartis, Basel, Switzerland), and vincristine (Hospira Worldwide Inc., Lake Forest, IL) (Sigma-Aldrich, St. Louis, MO) were dissolved in DMSO and stored at −20°C.

### Fluorescence Activated Cell Sorting (FACS)

For cell surface staining, cells were washed 1x with PBS and passed through a 40 μm filter (BD Biosciences, San Jose, CA) and blocked for 20 min at room temperature with FcR blocking reagent (Miltenyi Biotech, San Diego, CA). Cells were then stained with 1 µg antibody per 1×10^6^ cells for 15 min at room temperature in the dark. After 2x washing with PBS, cells were resuspended in 200 μL FACS buffer (1x PBS, 1% FBS, 1 mM EDTA, 25 mM HEPES pH 7) and analyzed on a BD Accuri C6 cytometer (BD Biosciences, San Jose, CA). As a control, cells were stained with a fluorophore-conjugated mouse IgG control. For intracellular staining, cells were first fixed with IC Fixation buffer (eBioscience, San Diego, CA) for 30 min on ice, and permeabilized with Perm/Wash buffer (#554723, BD Biosciences) before blocking and staining.

### Cell cycle and apoptosis

For cell cycle analysis, cells were fixed and permeabilized prior to resuspension in PI/RNase Staining buffer (BD Biosciences) for DNA content measurements and FACS analysis. For apoptosis detection, cells were resuspended in 200 μL 1x binding buffer with or without 5 μL Annexin V (BD Biosciences, Billerica, MA). After 15 min incubation, cells were washed with PBS, resuspended in 500 μL FACS buffer and 5 μL propidium iodide (BD Biosciences) and analyzed by FACS. PHA-739358 (Danusertib) is a pan-Aurora kinases inhibitor with activity against all Aurora kinase family members (A, B and C). US7 treated with 1μM of PHA-739358 (24 hours) as positive control for cell cycle analysis. Cells fixed, permeabalized and stained with PI/RNase before FACS analysis.PHA-739358-treated cells present an increased G2/M-phase population. Data were analyzed using FCS Express 5 Plus Research Edition and FlowJo V10.

### Viability and proliferation, drug treatment

Leukemia cells were cultured with an irradiated OP9 stroma layer in 24-well plates in complete αMEM in the presence of control DMSO or drugs (nilotinib, vincristine, Cpd 14 or Cpd17). Except where indicated otherwise, viable and total cell numbers were determined using Trypan blue exclusion. Growth curve assays were done by counting live cells using Trypan blue exclusion and an inverted microscope for >12 days after plating the cells during which fresh media was supplied every 2-3 days. In long-term assays, fresh drugs were faded with the medium changes. Cpd14-induced inhibition of cell growth was also measured using a colorimetric AlamarBlue cell viability reagent (#PI88951, Thermo Fisher Scientific, Waltham, MA). Cells (0.5-1×10^5^) were plated in 100 μL final volume in a 96-well plate in the presence of the different concentrations of Cpd14 for 24 hours at 37°C. Then 10 μL of the AlamarBlue reagent was added to each well. Within 5-6 hours incubation at 37°C, the fluorescence intensity at 535 and 590 nm was measured. Cell viability was calculated with respect to the control samples and reference background wavelength. At least three independent experiments were performed.

### Adhesion

A 96-well plate was coated with 5 μg/mL fibronectin in PBS, overnight at 4°C for 20 min at room temperature. BCP-ALL wells were washed with 0.1% BSA in PBS prior to blocking with 2% BSA in PBS for 1 hour at room temperature. Wells were then washed twice with 0.1% BSA in PBS and labeled with 5 µM Calcein AM (#C3100MP, Thermo Fisher Scientific, Waltham, MA) for 30 min at 37°C, then washed 2x with PBS prior to seeding at 5×10^4^ cells/well in αMEM base media with or without drugs for 30 min. β-Lactose (Sigma-Aldrich, St. Louis, MO) was included as a positive control for galectin-3 binding inhibition. After 30 min, the plate was read at 485 nm on a Synergy HTX Multi-Mode Reader (BioTek, Winooski, VT). The plate was washed two more times with 0.1% BSA in PBS to completely remove unbound cells and read again at 485 nm. The percent adherent cells was calculated using the following formula: [(RFU post wash-RFU background)/ (RFU pre wash-RFU background)] ×100%. To measure US7 and TXL2 adhesion to OP9 EV and OP9-KO cells, we collected floating cells and, after trypsinization, OP9 and associated ALL leukemia cells. Trypan blue was used to identify living cells, and manual counting was used to identify BCP-ALL cells in the mixture samples.

### Migration

Migration of BCP-ALL cells was also tested using a 24-well Transwell system (5 µm pore size) (#3421, Costar, Corning Incorporated, NY). Migration assays were carried out in αMEM with 2% FBS. BCP-ALL cells (0.5-1×10^5^) were plated inside the upper mesh insert (100 µL volume) while either 200 ng/mL SDF-1α (#300-28A, PeproTech, Rocky Hill, NJ) or a monolayer of OP9 cells (1-2×10^4^ cells/well) in the lower chamber (final 600 µL of media) served as chemoattractant. The number of migrated BCP-ALL cells in the lower chamber was counted by Trypan blue exclusion, with migration to OP9 measured after overnight incubation, and migration to SDF-1α after 2-4 hours incubation.

### Plasmids and CRISPR/Cas9

Galectin-3 cDNA was subcloned into the pMIG-GFP retroviral expression vector (plasmid #9044, AddGene, Cambridge, MA). Stably transduced hGalectin-3 expressing OP9 cells were generated by infecting cells with retroviral particles and FACS sorting of GFP expressing (positively transduced) cells [11]. Western blotting after cell expansion was used to confirm successful galectin-3 expression.

To induce CRISPR/Cas9-mediated deletion of Gal3, the LentiCRISPR v2 plasmid containing Cas9 and including a puromycin selection marker was purchased from Addgene (Plasmid #52961). A 20 nucleotide sgRNA sequence targeting exon 3 of mouse *lgals3* was designed using the website http://crispr.mit.edu/. The complementary sgRNA oligonucleotides (forward): 5’-caccgTCAAGGATATCCGGGTGCAT-3’ and (reverse): 5’-AAACATGCACCCGGATATCCTTGAC -3’, (synthesized by Integrated DNA Technology) were annealed and cloned into the LentiCRISPR v2 plasmid, and DNA sequences were verified. Clones were first verified by restriction digestion with BsmBI.

CRISPR-induced mutations were detected using a Surveyor Mutation Detection Kit (#706025, Integrated DNA Technologies) according to the manufacturer’s instructions. Frequency and type of mutations were determined by cloning the OP9 genomic DNA fragment spanning 600 bp around the gRNA DNA break site into pUC19 plasmid (#50005, AddGene) and sequencing 50 single colonies (primers: (forward): 5’-AGGCCAGAACAAGACATGATACA-3’ and (reverse): 5’-ACCAATGTCCCCTCCACTTG-3’). Serial dilutions in a 96-well plate were used to isolate and identify single colonies with bi-allelic homozygous mutations. Western blotting was done to detect complete Gal3 expression knockout. CRISPR/Cas9 mediated Lgals3 homozygous disruption occurred at frequencies of around 32% and resulted in three genotypes (Fig S2).

### Western immunoblot

Whole cell lysates were prepared by suspending cell pellets (on ice) for 20 min in RIPA buffer (1% NP-40, 0.1% SDS, 150 mM NaCl, 50 mM Tris, pH 8.0) supplemented with 1x complete EDTA-free protease inhibitor and 1x PhosStop (Roche, Basel, Switzerland). Lysates were separated by 4-20% SDS-PAGE and transferred to PVDF membranes (GE Healthcare, Piscataway, NJ). Membranes were blocked with 5% BSA in 1x TBS with 0.1% Tween 20 for 1 hour at room temperature with shaking. Primary antibodies for actin (sc-47778, Santa Cruz Biotechnology, Santa Cruz CA), Galectin-1 (GTX101566) (GeneTex, Irvine, CA), and galectin-3 (#125402, Biolegend, San Diego, CA), were diluted 1:1000 or 1:2000 in 5% BSA and incubated for 1hour at room temperature. Membranes were washed using 1x TBS-T and incubated with HRP-conjugated secondary antibodies diluted in 1x TBS-T with 5% BSA, washed, and developed using SuperSignal West Dura Chemiluminescent Substrate (#32106, Thermo Fisher Scientific).

### Statistical analysis

Biological experiments were analyzed with paired or unpaired t-tests and ANOVA using GraphPad Prism and Excel software. The value of p < 0.05 was considered to be statistically significant. Treatments were carried out in triplicates and at least in two independent experiments.

## Results

### Bone marrow stromal cells as source of galectin-3

Pediatric patients with BCP-ALL are typically treated with a 1-month regimen of induction chemotherapy of which vincristine is usually one component. Thereafter, patient BM is examined for the presence of minimal residual disease (MRD). The absence of MRD at this time point is highly correlated with event-free and overall survival in both pediatric and adult BCP-ALL [22]. As shown in **Fig. S1A**, the reduction in CD19 and CD10 mRNA expression as hallmarks of BCP-ALL cells in these BM specimen reflects the massive eradication of the leukemia cells. Interestingly, there is a marked increase in galectin-3 mRNA in the same samples, suggesting that chemotherapeutic treatment of the BM, including leukemia cells and cells in the leukemia microenvironment, induces high amounts of galectin-3. Thus, at the end of induction therapy, the BM is likely to contain high levels of galectin-3. Higher levels of galectin-3 (*LGALS3*) mRNA also correlate with a higher probability of being MRD-positive at the end of such induction therapy (**Fig. S1B**). Moreover, when patients relapse, samples show increased *LGAL3* mRNA compared to diagnosis samples from the same patient (**Fig. S1C**).

The bone marrow tumor niche where BCP-ALL develops consists of different stromal cell types, some of which contribute in a major way to the survival and growth of the leukemia. The mesenchymal stromal cells (MSC) are such key components [23, 24]. To determine if these cells make galectin-3 we grew MSC out from primary human bone marrows. FACS analysis of CD45^neg^, SSEA4^pos^, CD271^pos^ confirmed that this entire cell population is positive for Gal3 expression (**Fig. 1A**, MSC-1 and MSC-2). We also compared the ability of such MSC to support BCP-ALL cells next to a human MSC cell line and a murine bone marrow MSC cell line OP9. Based on viability, all three MSC were able to keep the BCP-ALL cells alive in such co-culture systems ((**Fig. 1B, Fig. 1C, left panel**). However, interestingly, the OP9 cells allowed more cell proliferation compared to the MSC and were also superior in providing protection against chemotherapy treatment with vincristine (**Fig. 1C, right panel**). Because OP9 cells allow better BCP-ALL growth, have a longer life span than primary MSC and are more consistent in their own growth, we used these cells for subsequent experiments.

**Figure 1.**
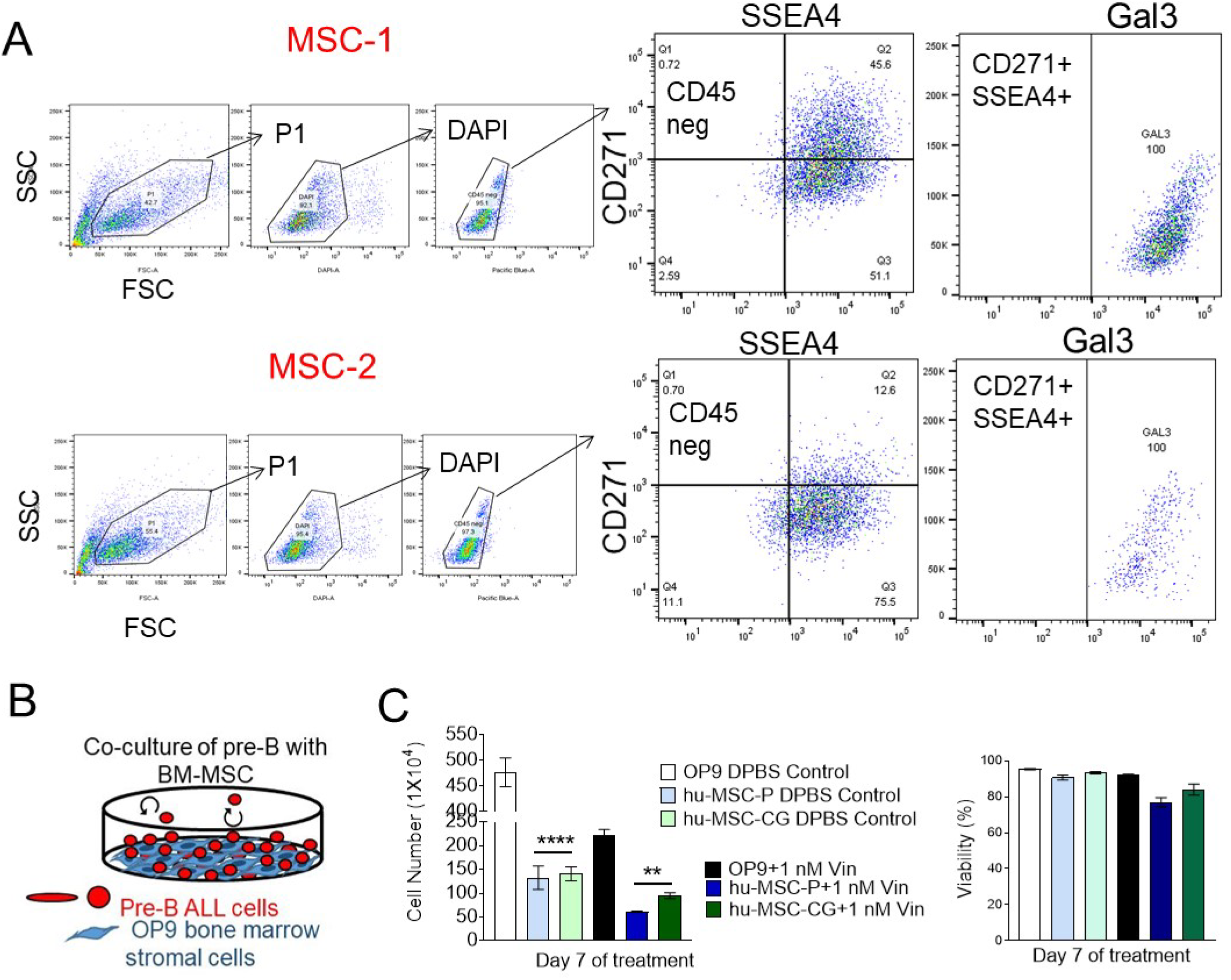
Bone marrow stromal cells produce Galectin-3. (A) MSC from two different normal BMT donor screens MSC-1 and MSC-2 were grown out as adherent cells. FACS analysis on the indicated populations using the gating strategy is shown in the figure.(B) schematic of co-culture system of leukemia cells with stromal layer. Stromal cells have been mitotically inactivated. They support the leukemia cells but no longer can divide. (C). Comparison of human primary bone marrow MSC with mouse bone marrow MSC OP9 cells for ability to protect human BCP-ALL US7 cells against chemotherapy. Left panel: viability (viable cell number/total cell number x 100) determined using Trypan blue. Right panel: total cell number Stromal cells were mitotically inactivated by treatment with 10 μg/ml mitomycin C. Analysis on day 7 of 1 nM vincristine treatment (comparison with OP9 cells, triplicate samples. One-way ANOVA, multiple comparisons). hMSC-1 and hMSC-CG, primary and immortalized human MSC. **p<0.01, ***p<0.001

### Gal3 made by stroma is a major source for BCP-ALL cells-

We then examined Gal3 expression in a primary BCP-ALL sample by FACS (**Fig. 2A)**. Initially the sample had a signal for Gal3 that was higher than normal BM. After a 10-day co-culture with wild type OP9 cells the signal was clearly increased, suggesting uptake of stromal-produced gal3 in these cells. To examine this, we used CRISPR/Cas9 to knock out Gal3 in OP9 stroma cells (strategy in **Fig. S2**). We used single-cell cloning to isolate clones with bi-allelic Gal3 knockout (**Fig. S2C**). Compared to empty vector-transduced OP9 (OP9-EV), the OP9-Gal3-KO cells had a minimal signal using FACS (**Fig. 2B**). We then used the two different OP9 genotypes (KD and EV) for co-culture with two different human BCP-ALLs, US7 and TXL2. As shown in **Fig. 2C**, when these leukemias were harvested from above the OP9-Gal3-KO cells, no Gal3 was detected by Western blot. In contrast, when the same cells were co-cultured with OP9-EV cells, Gal3 protein was detected. These results are consistent with our previous report in which we found Gal3 uptake by BCP-ALL cells [12].

**Figure 2.**
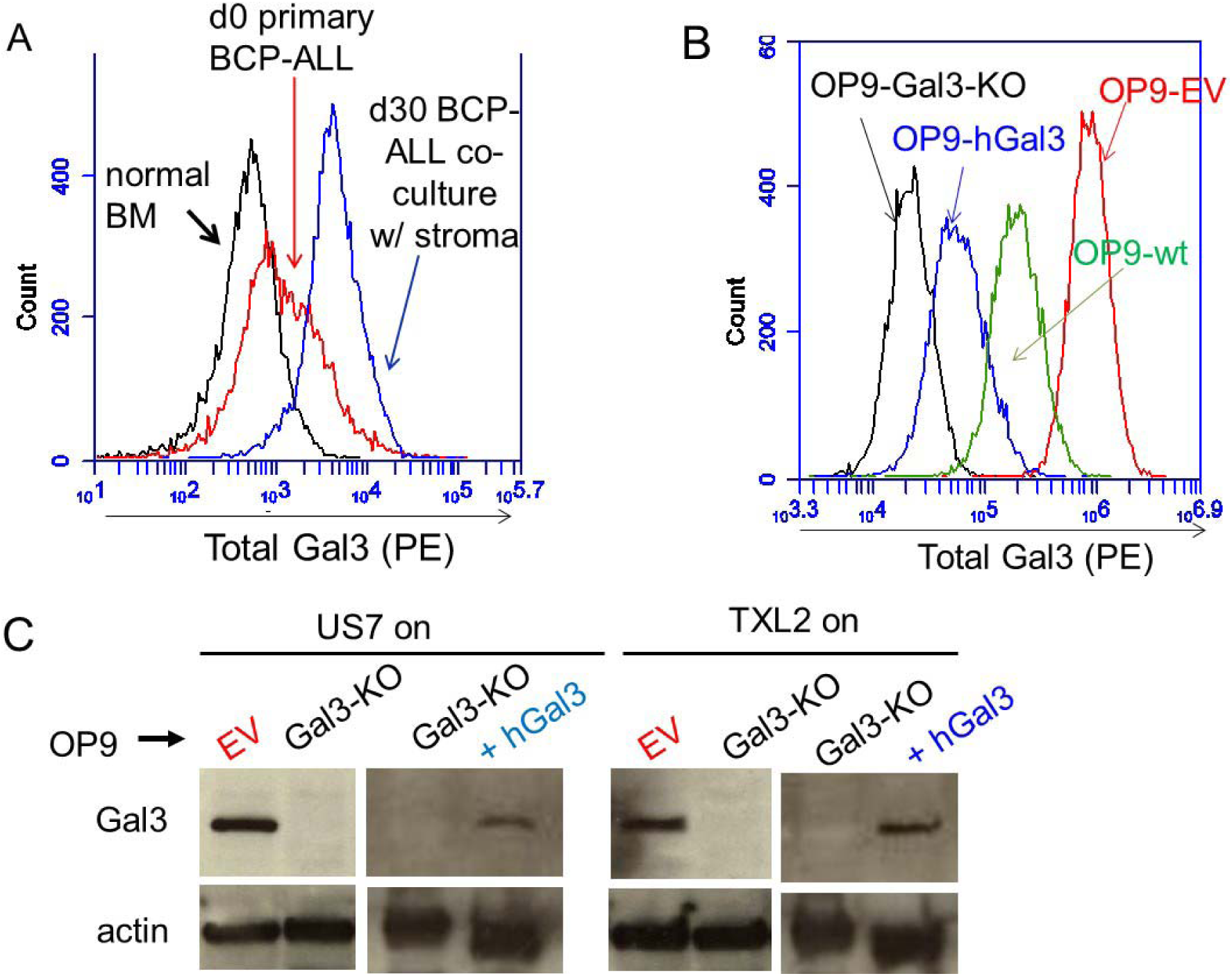
Protective stromal cells provide Gal3 to BCP-ALL cells. (A) FACS analysis of total Gal3 expression in primary leukemia sample LAX39 (Ph-, >90% CD10/CD19^+^). Gal3 expression in the BCP-ALL patient sample compared to normal bone marrow on day 0 and after ∼10 days co-culture with irradiated OP9 stroma cells. (B) Total Gal3 expression in 10,000 fixed and permeabilized OP9 stromal cells of the indicated genotypes. (C) Western blot of the indicated BCP-ALLs grown for >4 days with OP9-EV, Gal3-KO or Gal3-KO + hGal3 cells. BCP-ALL cells were harvested from the medium.

### Effect of Gal3 loss on BCP-ALL homeostasis

We first examined the effects of loss of stromal-produced Gal3 on BCP-ALL cell viability and proliferation. Both US7 and TXL2 grew normally *in vitro* when co-cultured with both Gal3-KO and control OP9 cells over a two-week co-culture: viability (**Fig. S3A**) and proliferation (**Fig. S3B**) in the absence of stress were normal and accordingly, no change was detected in the BCP-ALL cell cycle as determined by FACS-assisted measurement of DNA content (**Fig. S3C)**. These observations indicate that, under steady-state conditions, stromal Gal3 expression does not regulate BCP-ALL cell survival and proliferation.

BCP-ALL cells depend on supportive stromal cells for sustenance and expansion in culture in part because they migrate towards specific chemokines secreted by these cells and then adhere to and migrate underneath the stroma. Because Gal3 has been implicated in regulating both normal and tumor cell motility, we examined if extracellular Gal3 levels can also modulate BCP-ALL movement. To measure the association of the leukemia cells to Gal3-deficient stroma, we determined the number of living floating and adhered BCP-ALL cells via Trypan blue exclusion. This assay showed that significantly lower numbers of BCP-ALL cells were found to be adhered to OP9-Gal3-KO cells than to control OP9 stroma after a 24 or 72 hours co-culture and conversely, that more cells remained floating in the medium (**Fig. 3A**, US7; **Fig. 3B**, TXL2). To demonstrate that this was Gal3-dependent, we also reconstituted the OP9-Gal3-KO cells with human Gal3. Although expression levels were lower than that of WT OP9 (**Fig. 2A**) it was sufficient to restore association with the OP9 cells (**Fig. 3C**). We also performed a Transwell migration assay and found that fewer cells migrated down to OP9 cells with no Gal3 production than to control OP9 cells (**Fig. 3D**). These findings point to a key role for stromal-produced Gal3 in mediating the migration and adhesion of BCP-ALL cells to the crucial protective stromal components of the tumor microenvironment.

**Figure 3.**
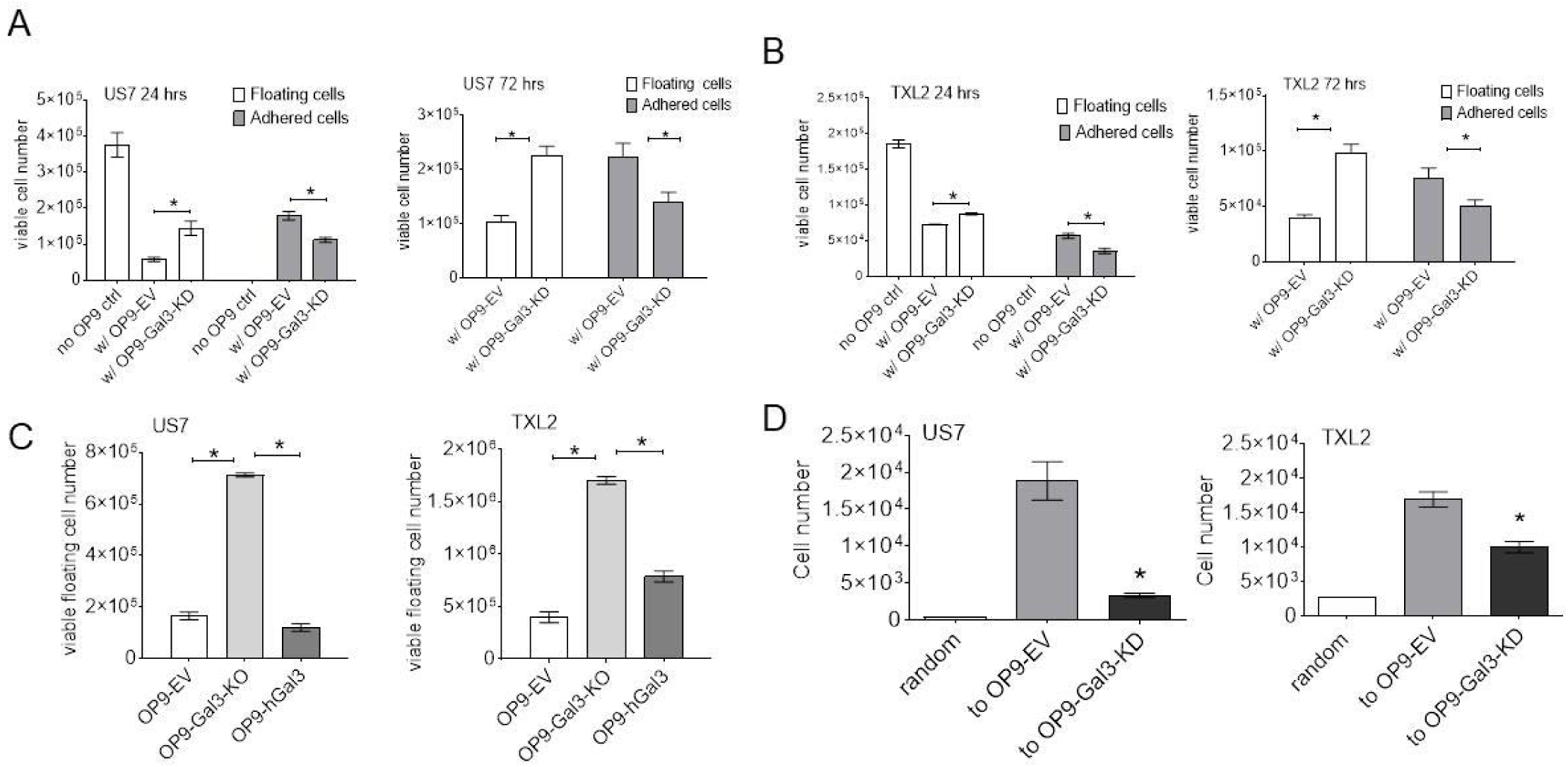
Gal3 deficiency impairs BCP-ALL cell adhesion and migration. Adhesion of BCP-ALL cells to Gal3-deficient OP9 and control stroma presented as number of viable cells. (A, B) BCP-ALL cells in the supernatant (floating) and attached to OP9 stroma (adhered) were both determined. (A) US7 cells after 24 and 72 hours of migration (B) TXL2 cells at 24 and 72 hours. (C) Adhesion of US7 or TXL2 cells to the OP9 cells of the indicated genotype measured after 24 hours. OP9-hGal3: OP9 Gal3-KO cells expressing hGal3. (D) Migration of US7 or TXL2 cells over a 4 hour period toward OP9 Gal3-KO stroma and control OP9-EV cells measured using a Transwell assay. Motility in the presence of medium was used as readout for random (spontaneous) migration. Error bars, mean± SEM of duplicate (C-D) or triplicate (A-B) samples. * p < 0.05, t-test.

### Depletion of Gal3 increases BCP-ALL cells response to drug treatment

We next examined if stromal-produced Gal3 regulates drug sensitivity of BCP-ALL cells. BCP-ALL cells were cultured with either OP9-Gal3-KO or OP9-EV control stroma in the presence of 5 nM vincristine, part of the standard cytotoxic therapy in BCP-ALL (Vin, to treat Ph-US7 cells) or 20 nM of the targeted tyrosine kinase inhibitor nilotinib (Nilo, to treat Ph+ TXL2 cells) for >15 days. Cell viability was measured every 3 days during drug treatment and is presented as percent of total. Interestingly, as shown in **Fig. 4A, B**, deficiency of Gal3 in BCP-ALL cells increased the sensitivity of the BCP-ALL cells to the chemotherapy when compared to OP9-EV control. In both treatment groups, following a sharp decline, cell viability recovered around day 5 (**Fig. 4A, B**). However, viability of leukemia cells after day 5 was significantly lower (ANOVA of all time points US7: p=0.0009 and TXL2: p=0.028) in BCP-ALL cells expanded with Gal3-deficient stroma when compared to control OP9, suggesting that extracellular Gal3 can provide considerable chemoprotection for BCP-ALL cells. As expected, viability of BCP-ALL cells in the absence of chemotherapy (control DMSO treated co-cultures) remained unaffected regardless of the Gal3 status of the stroma.

**Figure 4.**
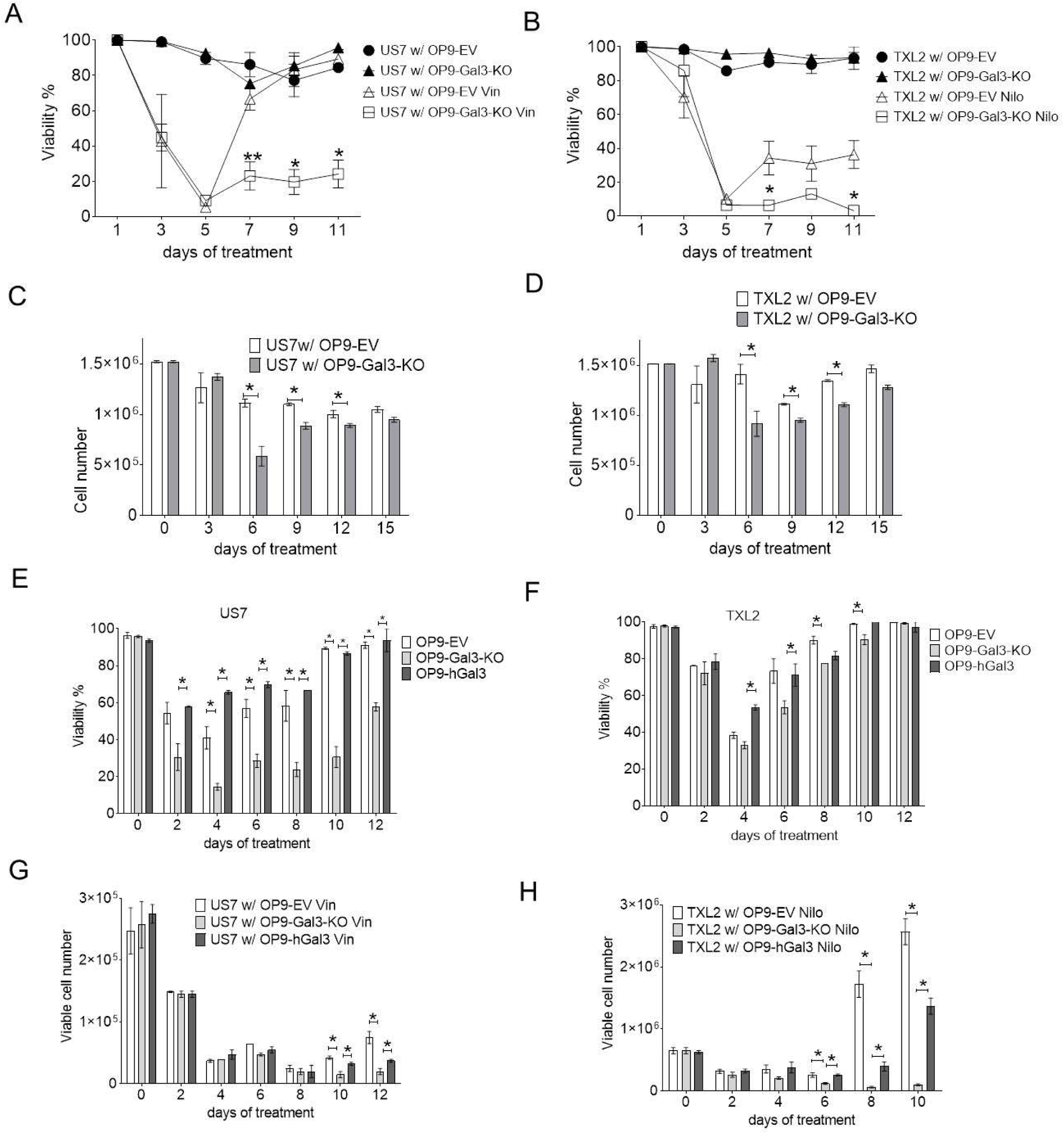
Depletion of Gal3 significantly enhances BCP-ALL cell response to chemotherapy. US7 cells (A, C, E, G) or TXL2 (B, D, F, G) were grown on OP9 cells as indicated. Cells were treated with solvent as control, or with 2 nM vincristine (vin; US7, A, C, E, G) or 20 nM nilotinib (nilo; TXL2, B, C, F, H). A-B and E-F viability [percentage of Trypan blue excluding cells/total cells] and C-D and G-H cell numbers. (E-G). Analysis similar to A-D, but also including OP9-Gal3 KO cells reconstituted with human Gal3 [hGal3]. Error bars, mean ±SEM of 4 replicates per time point (A-C) or 2 replicates per time point (E-F). * p < 0.05, **p<0.01.

In addition, leukemia cell proliferation (**Fig. 4C, D**) resumed after a few days of drug treatment and, similar to viability, BCP-ALL cells co-cultured with Gal3-deficient stroma had a significantly inferior (ANOVA of all time points US7: p=0.0001 and TXL2: p=0.0005) recovery in proliferation, suggesting that BCP-ALL cells lacking stromal-produced Gal3 responded better to chemotherapy. The expression of human Gal3 in the OP9 Gal-3 deficient cells was able to partly rescue both viability and proliferation (**Fig. 4E-F** and **G-H**, respectively). We conclude that Gal3 contributes to the microenvironment-mediated support against conventional drug treatment in BCP-ALL cells, and that inhibiting Gal3 may enhance BCP-ALL cell sensitivity to standard chemotherapy agents.

### Novel galectin-3 inhibitors-effect on BCP-ALL cell function

The design of specific small molecule inhibitors of Gal3 has been challenging because of the high amino acid sequence homology between members of the galectin family as well as the weak binding of carbohydrates and lectins, and low bioavailability [29]. We recently reported the development of a series of taloside-based compounds [21], Cpd14-Cpd18, with significant specificity for Gal3. The lead compound Cpd14 was able to significantly reduce viability in US7 and TXL2 in a 24 hour assay at 250 μM. We next tested the effects of Cpd14 and Cpd17 on important Gal3-regulated activities. BCP-ALL cells adhere to fibronectin (FN) in an integrin-dependent manner [25, 26]. As shown in **Fig 5A, B**, both US7 and TXL2 cells can adhere to fibronectin-coated plates ([controls). We compared the effect of Cpd14 on FN-mediated adhesion of the BCP-ALLs to that of β-lactose which is a natural competitor of Gal3 in binding to carbohydrate structures. Although Cpd14 is a monosaccharide-based compound, it was able to inhibit cell adhesion of TXL2 at an approximately 12-fold lower concentration than the disaccharide lactose (**Fig. 5B**). Interestingly, Cpd14 had a more potent inhibitory effect on migration of both US7 and TXL2 cells. Migration to both OP9 cells as well as to the chemokine SDF1-α secreted by stromal cells (**Fig 5C, D**). The IC_50_ of Cpd 17 was significantly lower than Cpd14 (83 μM and 307 μM) [27]. As shown in **Fig. 5F**, in agreement with this much lower amounts were needed to inhibit adhesion of BCP-ALL cells to OP9 stroma.

**Figure 5.**
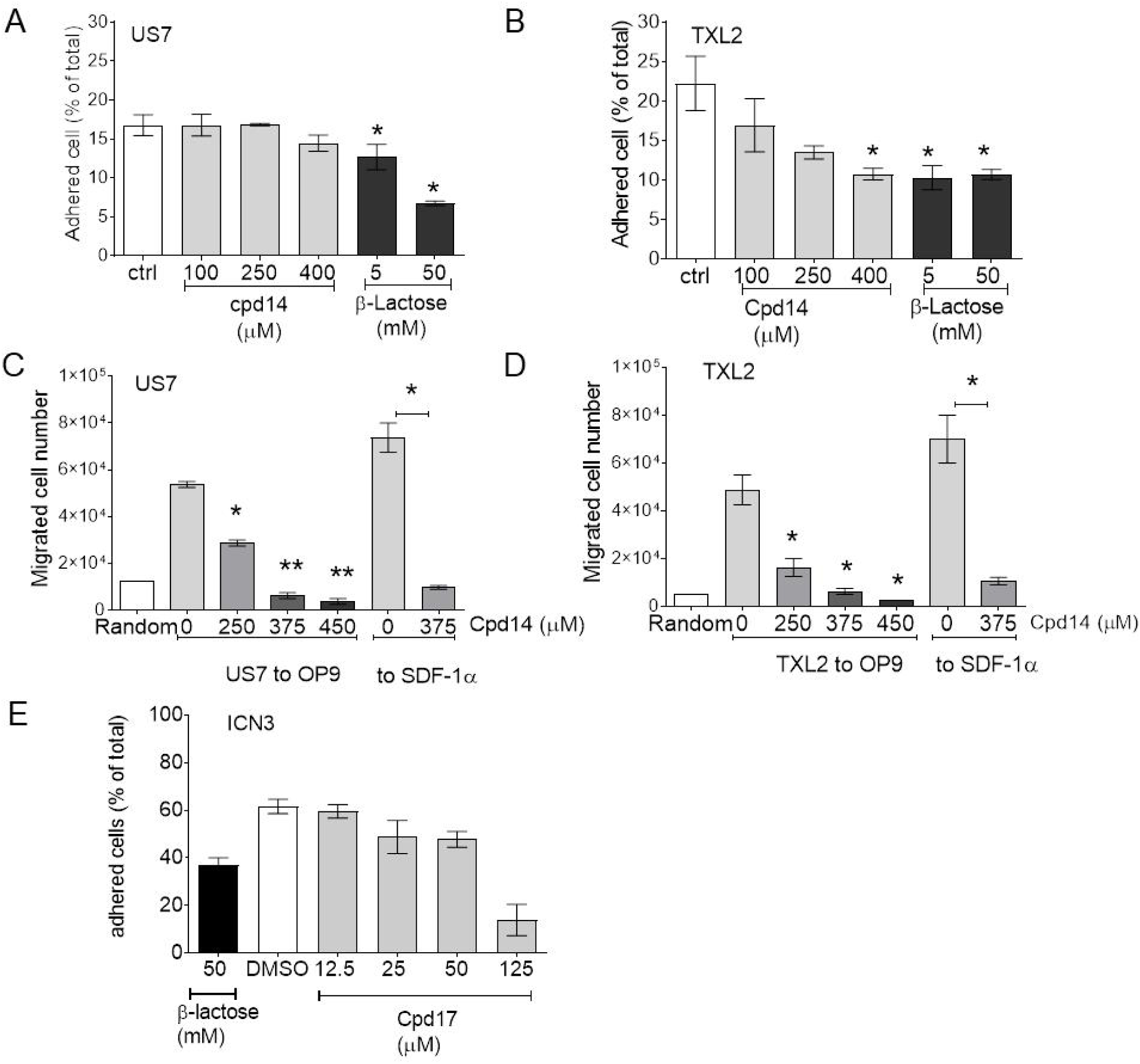
Effects of Cpd14 or 17 on BCP-ALL migration and adhesion. (A-B) Adhesion of US7 or TXL2 cells to fibronectin-coated wells with and without the indicated amounts of Cpd14 in a 30-min assay. (C-D) Migration of US7 or TXL2 to OP9 or to the chemokine 200 ng/mL SDF1-α (24 and 4 hours, respectively) in a Transwell assay when treated with the indicated concentrations of Cpd14. (E) Adhesion of ICN13 cells after 24 hours to OP9 in the presence of the indicated concentrations of Cpd17. Cells in suspension and adherent to (above and underneath) the OP9 cells were harvested and counted by Trypan blue exclusion. Results are presented as percentage of living [Trypan-blue excluding] cells adhering to OP9. Error bars, mean ±SEM of triplicate values (A, B, E) or duplicate wells (C, D) *p<0.05; **p<0.01. ctrl, DMSO control.

Because both Cpd14 and Cpd17 were able to inhibit BCP-ALL cell proliferation we also tested their effects on cell cycle. As summarized in **Fig. 6A**, the most clear effect of Cpd14 was to reduce the number of cells in S-phase (**Fig. 6B**, arrow). Compared to DMSO-treated cells, Cpd17 also reduced the percentage of cells in S phase, increasing those in G1. However, compared to the pan-Aurora kinase inhibitor PHA-739358, which also reduces the % of cells in S-phase but causes accumulation in G2, this effect was small (**Fig. 6C**). To investigate if BCP-ALL cells lose viability because of apoptosis we measured AV/PI expression using FACS. Compared to DMSO-treated controls, a 48 hour treatment with 250 or 500 μM Cpd14 caused a prominent increase in AnnexinV/PI^pos^ US7 or TXL2 cells [not shown]. This also applied to Cpd17 but at lower concentrations; a 24 hour treatment with 20 μM Cpd17 clearly increased the number of AnnexinV/PI^pos^ US7 cells (**Fig. 6D**).

**Figure 6.**
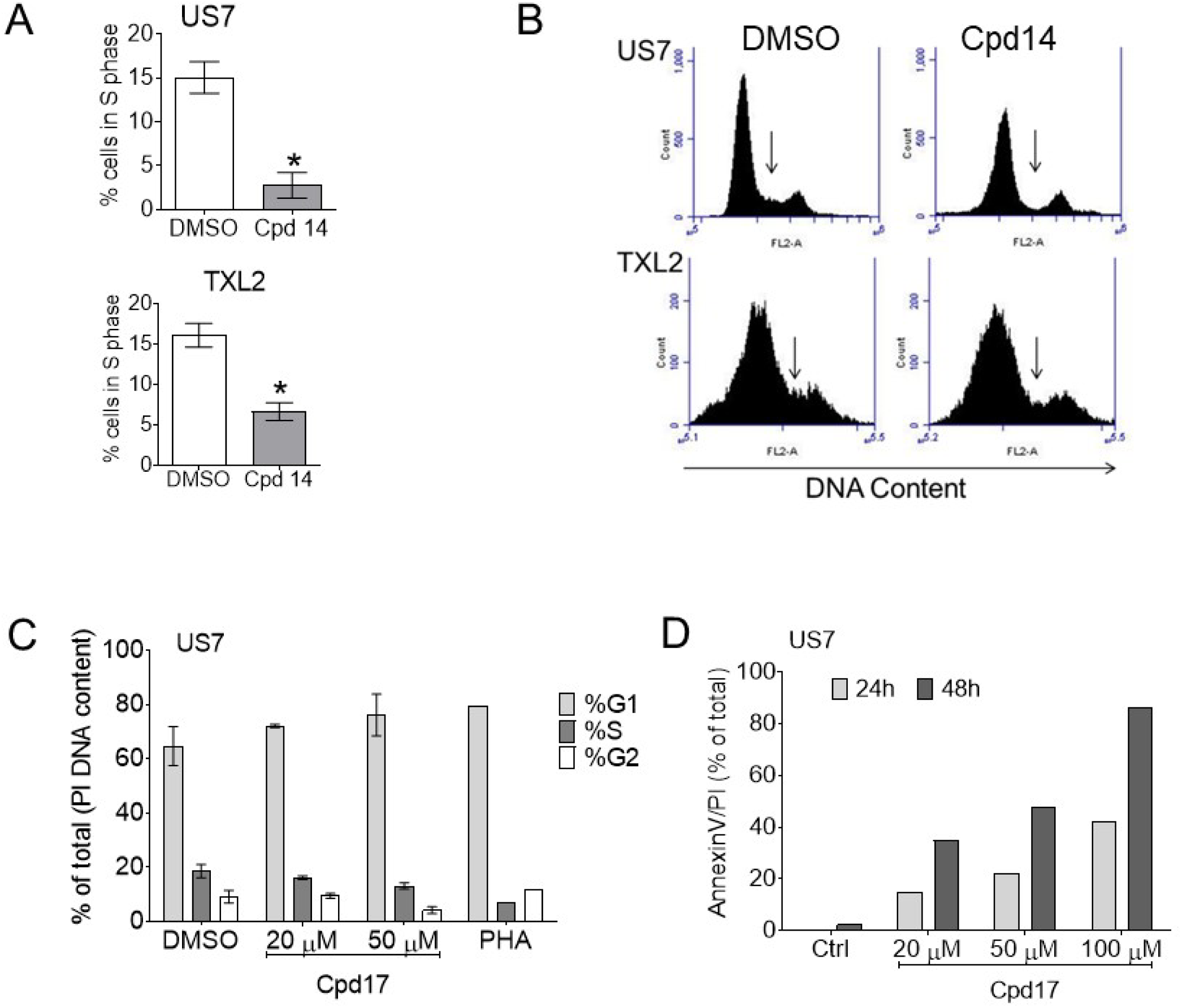
Analysis of effects of Cpd14 and 17 on cell cycle and apoptosis. (A-B): Cell cycle of US7 or TXL2 cells measured by DNA PI staining using FACS, following 48 hours exposure to 250 µM Cpd14. (A) Percentage of BCP-ALL cells in S phase; (B) representative image, arrow points to S phase DNA content. Error bars, mean ±SEM of duplicate measurements. * p < 0.05 (C-D) Treatment of US7 cells with the indicated amounts of Cpd17. (C) Cell cycle analysis using FACS and PI DNA staining. Duplicate measurements, differences not significant. PHA-739358 single sample. (D) percentage of apoptotic cells based on AnnexinV/PI positivity. PHA, 1 μM PHA-739358 (danusertib).

### Chemotherapeutic treatment of BCP-ALL combined with Cpd17

We next determined the IC_50_ for Cpd17 in five different human ALLs including TXL2 (diagnosis, Ph-positive), US7 (diagnosis, no karyotype abnormalities), ICN06 (ETV6-Runx1), ICN3 (MLL-AF4, infant ALL) and ICN13 (MLL-AF4, infant ALL). As shown in **Fig. 7A**, different BCP-ALLs had similar IC_50_ values for Cpd17. When tested over an 8-day period in the presence of OP9 cells (**Fig. 7B-E**, top, cell counts; bottom, viability), Cpd17 used at approximate IC_50_ value suppressed BCP-ALL cell growth by 50 % on day 3, which is in consistent with results obtained with Alamar Blue in **Fig. 7A**. Cpd17 continued to suppress BCP-ALL cell growth after day 3 (**Fig. 7B-E** top) and thus clearly, the compound is cytostatic at this concentration. In addition, viability of the cells decreased starting at day 3 (**Fig. 7B-E** bottom panels), indicating the compound is cytotoxic at this concentration.

**Figure 7.**
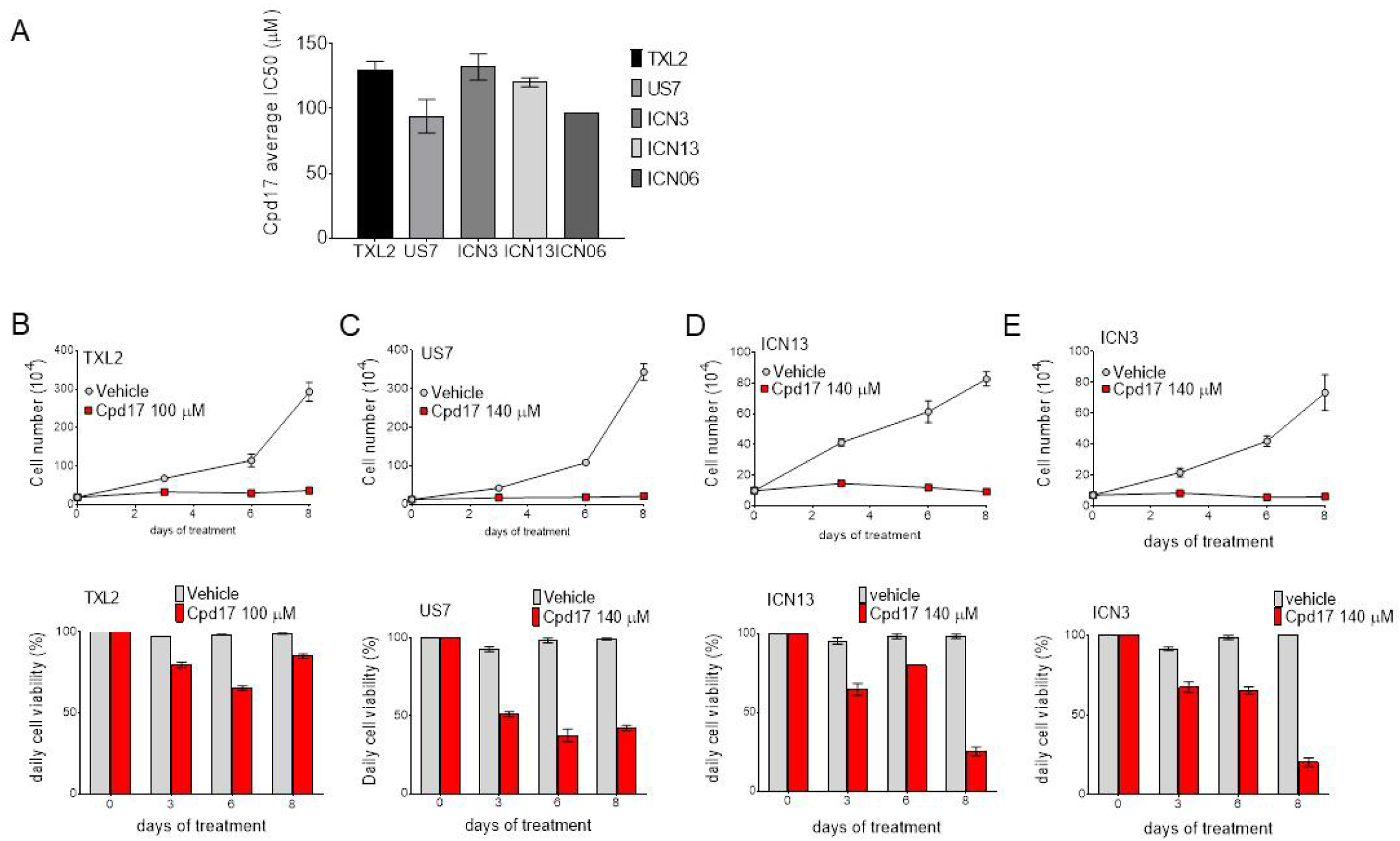
Determination of IC_50_ values for Cpd17. (A) For IC_50_ determination cells were incubated with different concentrations of Cpd17 for 3 days without OP9 cells. Cell counts were done using Alamar blue. Error bars, mean ±SEM of independent triplicate IC_50_ determinations. (B-E) Different ALLs as indicated were co-cultured with OP9 cells for 8 days and treated with ≈IC_50_ concentrations of Cpd17. Top panels, cell numbers; lower panels viability determined by Trypan blue exclusion. Representative results from 3 independent experiments. Note: cell viability is defined here as the number of Trypan blue excluding cells in Cpd17 treatment group/number of trypan blue excluding cells in group treated with solvent DMSO at each time point. Error bars, mean ±SEM of triplicate samples.

Next, we tested the possibility of combination treatment with Cpd17 on a number of different BCP-ALL cells, measuring both cell counts (**Fig. 8**) and cell viability (**Fig. S4**) over a 23-day treatment of leukemia cells in co-culture with OP9 stroma to allow detection of the emergence of relapse. We first determined the lowest effective concentration of Cpd17 for a drug combination treatment of US7 cells. US7 cells were treated with 2 nM vincristine alone, and with either 1 or 5 µM Cpd17 alone, or a combination of vincristine and Cpd17, as indicated in the figure. As expected, (**Fig. 8A and Fig. 8B)**, Cpd17 at concentrations of 1 μM and 5 μM did not affect US7 cell growth compared to control (DMSO). Vincristine at 2 nM suppressed proliferation of the cells, but after around day 10, relapses emerged in that the cells resumed growth in spite of the continued presence of vincristine. Interestingly, in the combination treatment of 2 nM vincristine with 1 μM Cpd17, relapse was clearly suppressed although not eliminated (**Fig. 8A**). In contrast, when 5 μM Cpd17 was combined with 2 nM vincristine, cells were no longer able to divide, even after an extended period of observation (**Fig. 8B**).

**Figure 8.**
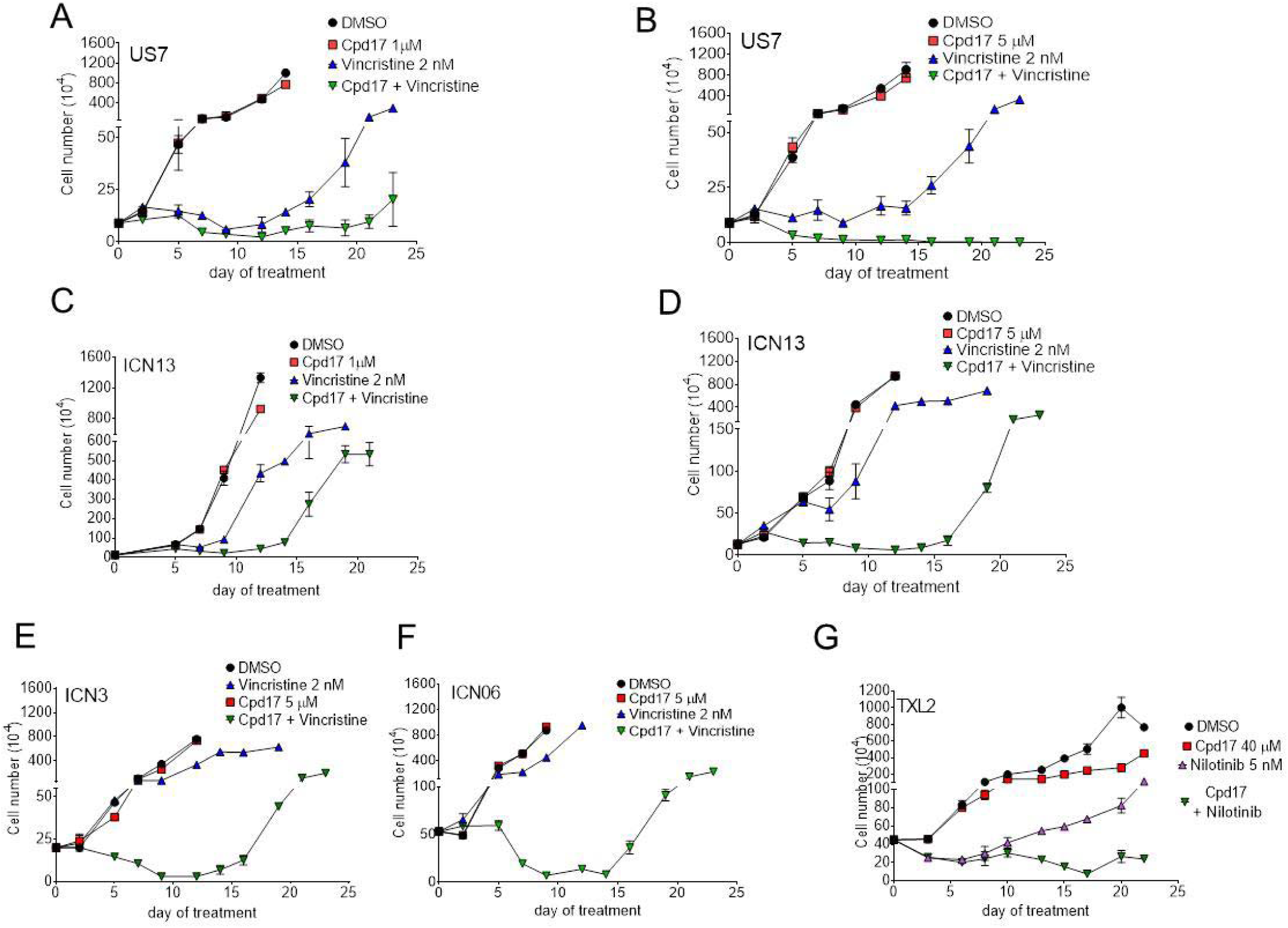
Proliferation of BCP-ALLs treated with combination treatment with Cpd17. Samples were treated with solvent DMSO [black], Cpd17 [red], standard chemotherapy [blue] or a combination of chemotherapy with Cpd17 [green]. Fresh drugs were added with each change of medium. Cell counts by trypan blue exclusion in DMSO and Cpd17 groups were terminated (on day 12-14 for DMSO and Cpd17-only samples, later in vincristine-only resistance) when growth exceeded well capacity in panels A-F. (G): Cpd17 was administered at 10 μM on day 0, increased to 20 μM on day 6 and further increased to 40 μM on day 10 as indicated by #, because drug combination effects were minimal at 10 or 20 μM. Results presented are average of three replicates +/- SD of one experiment. A-F: 2 nM vincristine as standard chemotherapy; G: 5 nM targeted tyrosine kinase inhibitor nilotinib. A-B: US7 cells; C-D: ICN13; E: ICN3; F: ICN06; G: TXL2. A, C: 1 μM Cpd17. B, D-F: 5 μM Cpd17. G: 40 μM Cpd17. Note the discontinuity of the Y-axis.

Similar to US7, the growth of ICN13 was not affected by 1 or 5 μM Cpd17 (**Fig. 8C, D**). Cell division of ICN13 was inhibited less than US7 by treatment with 2 nM vincristine. However, also with this BCP-ALL, the combination of Cpd17 with vincristine showed synergistic cytotoxic effects in ICN13 cells but relapse did emerge with the drug combination. Viability also dropped in the first 9-10 days, but cells become resistant.

ICN3 and ICN06 BCP-ALLs were largely vincristine-resistant (**Fig. 8E, F**, compare DMSO to 2 nM vincristine) and Cpd17 alone at 5 μM was not cytostatic either. Nonetheless, when 2 nM vincristine was combined with 5 µM Cpd17, growth was suppressed over at least 10 days of treatment. The viability of the cells also decreased in the first 9-10 days of combination treatment, after which resistance developed (**Fig. S4E, F**).

TXL2 BCP-ALL is driven by an oncogenic Bcr/Abl fusion protein and is treated with the targeted tyrosine kinase inhibitor nilotinib. Nilotinib on its own was not cytotoxic, but did inhibit the proliferation of these leukemia cells (**Fig. 8G**, compare DMSO and nilotinib). The addition of 40 µM Cpd17 to the nilotinib further reduced proliferation and modestly reduced viability of the cells. However, resistance to the combination treatment developed at day 20. Additive cytotoxic effects were seen at Cpd17 40 μM and nilotinib 5 nM from day 13 but the effects were reversed on day 20.

## Discussion

Gal3 has been implicated in numerous pathologies [15, 28], is present at different subcellular locations [29-34] and has almost 250 cell-surface binding partners [35]. Because of this, the exact mechanism by which stromal produced extracellular Gal3 protects BCP-ALL cells against the conventional chemotherapy drugs vincristine and nilotinib, as found in our study, is difficult to determine. However, survival and proliferation of BCP-ALL cells is strongly dependent on their ability to migrate and adhere to specifically protective microenvironmental niche cells in the bone marrow [36]. When such interactions are inhibited, for example by using AMD3100, next generation CXCR4 inhibitors or integrin-blocking strategies, BCP-ALL cells are mobilized and become easier to eradicate with conventional chemotherapy [5, 7, 26, 37-39]. We here show that extracellular Gal3 produced by stromal cells functions as one of the signals to BCP-ALL cells which regulates leukemic cell adhesion and migration. Therefore, we argue that one of the mechanisms through which reduction in extracellular Gal3 activity makes BCP-ALL cells more vulnerable to drug treatment is through an indirect route, by interfering with so-called cell adhesion-mediated drug resistance [40, 41].

How Gal3 produced by stromal cells can promote BCP-ALL migration is currently not clear, but it could cluster key glycoprotein binding partners such as integrins on the plasma membrane and promote cell polarization and subsequent directed migration. A second possibility is that Gal3 from stroma regulates intracellular events after its uptake by the BCP-ALL cells, as we have previously shown [12] and confirmed (result not shown). For example, in prostate cancer cells, Gal3, through stabilizing focal adhesion kinase at focal adhesions, can promote cancer cell motility [42]. In HeLa cells, Gal3-mediated activation of the Erk1/2 pathway was shown to be necessary for cell migration [13]. One intracellular mechanism through which Gal3 could polarize cells is through its interaction with Myh2a, an interaction previously reported by Nakajima et al [43]. By a recombinant Gal3 affinity column and confirmed through co-immunoprecipitation, we have also detected a direct protein-protein interaction between NMIIA (MYH9) and Gal3. Because NMIIA inhibition using blebbistatin resulted in a significant reduction in BCP-ALL adhesion to fibronectin, and migration toward SDF1α as well as OP9 cells, the intercellular interaction between these proteins could regulate these BCP-ALL activities (results not shown).

Pharmacologic targeting of Gal3 has been proposed as an attractive novel therapy for acute myeloid leukemia [44-47]. Our knock out experiments of Gal3 in the stromal cells provide a proof-of-principle that a reduction in Gal3 (activity) would also be beneficial in reducing the chance of persistence of MRD in the induction chemotherapy phase of treatment of BCP-ALL. However, the specific targeting of Gal3 using drugs or drug-like compounds has been a considerable challenge due to the highly conserved carbohydrate recognition domain shared by all galectins. Here, we tested Cpd14 and Cpd17 as novel, talopyranoside-based [21] small molecule antagonists in a number of relevant biological assays. Apart from the clear effect on cell migration, these compounds also had cytostatic and even cytotoxic effects when used alone. Because some of these assays were done in the absence of OP9 cells, all (inhibitable) Gal3 would have to be that which is endogenous to BCP-ALL cells. Levels of Gal3 are low in non-stressed BCP-ALL cells, but these cells do synthesize Gal3 *de novo* when subjected to stress [11, 12]. However we can not exclude the possibility of significant off-target effects, in particular because high amounts of compounds were needed. On the other hand, Galectins do have a weak binding affinity for their carbohydrate ligands and from this perspective Cpd17 as a monosaccharide-based compound did have an impressive activity on BCP-ALL cells. We conclude that these are promising compounds, but will require further optimization before they are suitable for *in vivo* preclinical studies. For example, we found that Cpd17 is not chemically stable and even in 100% DMSO stored at −20°C, its activity markedly diminished. We also were unable to test Cpd17 in animal experiments because it is very hydrophobic and it was not possible to find a biocompatible solvent.

By a careful titration of the drugs used in the chemotherapy combination experiments, we were able to show useful effects of Cpd17 at low micromolar (5 μM) amounts when combined with doses of vincristine that were unable to prevent BCP-ALL cell proliferation. This warrants further development of Cpd17 or derivatives to increase solubility. Since Gal3 is implicated in other malignancies, the impact of understanding how it regulates motility and invasion will not be limited to BCP-ALL. Importantly, our studies show that it is feasible to target the cross-talk between the tumor microenvironment and BCP-ALL cells via carbohydrate-lectin interactions, and show that Gal3 inhibition could be a novel approach to treat BCP-ALL.

## Acknowledgements

N.H. was supported by NIH PHS RO1 CA172040 and CA090321. The Cancer Council Queensland (ID1043716, ID1080845) is acknowledged for financial support (H.B, UJN). K.B was supported by Griffith University GUPRS and GUIPRS scholarships. We thank Charlotte Glackin for sharing the immortalized human MSC cell line. We acknowledge Dr. Haike Ghazarian for co-culture with human MSC, and the substantial contributions of Dr. Miao-Juei Huang in the detailed analysis of the effects of Cpd17 on BCP-ALL cells.

---

**Figure S1.**
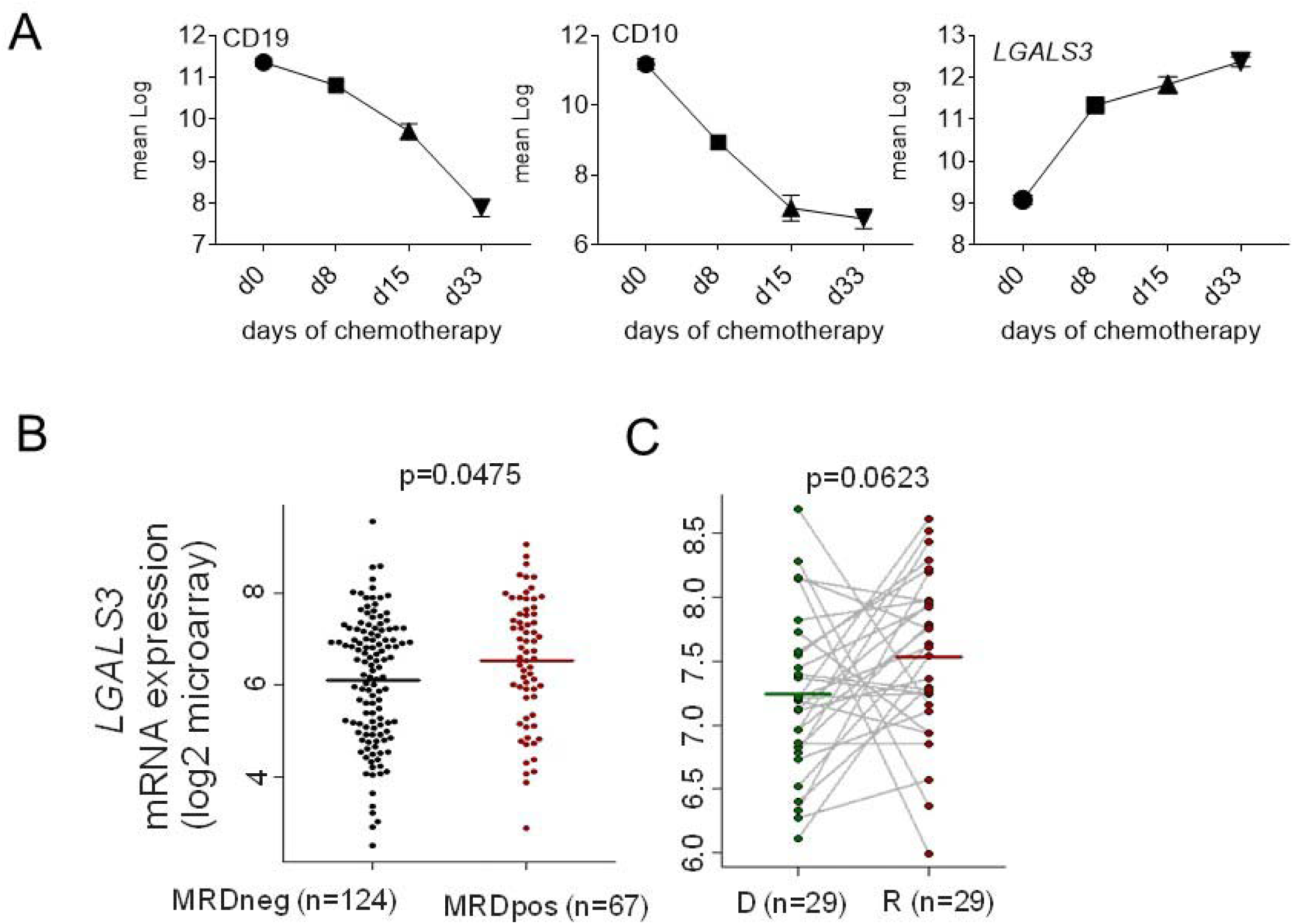
High levels of Galectin-3 mRNA in pediatric BCP-ALL. (A) Mean log-transformed normalized GEP values for the indicated genes on 220 pediatric de novo ALL at diagnosis, day 8, day 15, and day 33 of remission-induction therapy (48) (GSE67684). (B) *LGALS3* expression for probe set 1557197_a_at in the indicated samples [gene microarrays, log2 expression, COG P9906, GSE11877). p-value, two-sided Wilcoxon test. (C). *LGALS3* in paired early (<36 months) relapse and diagnosis pediatric ALL samples.(Gene microarray array expression (log2) COG P9906 GSE28460). p-values, paired two-sided Wilcoxon test.

**Figure S2.**
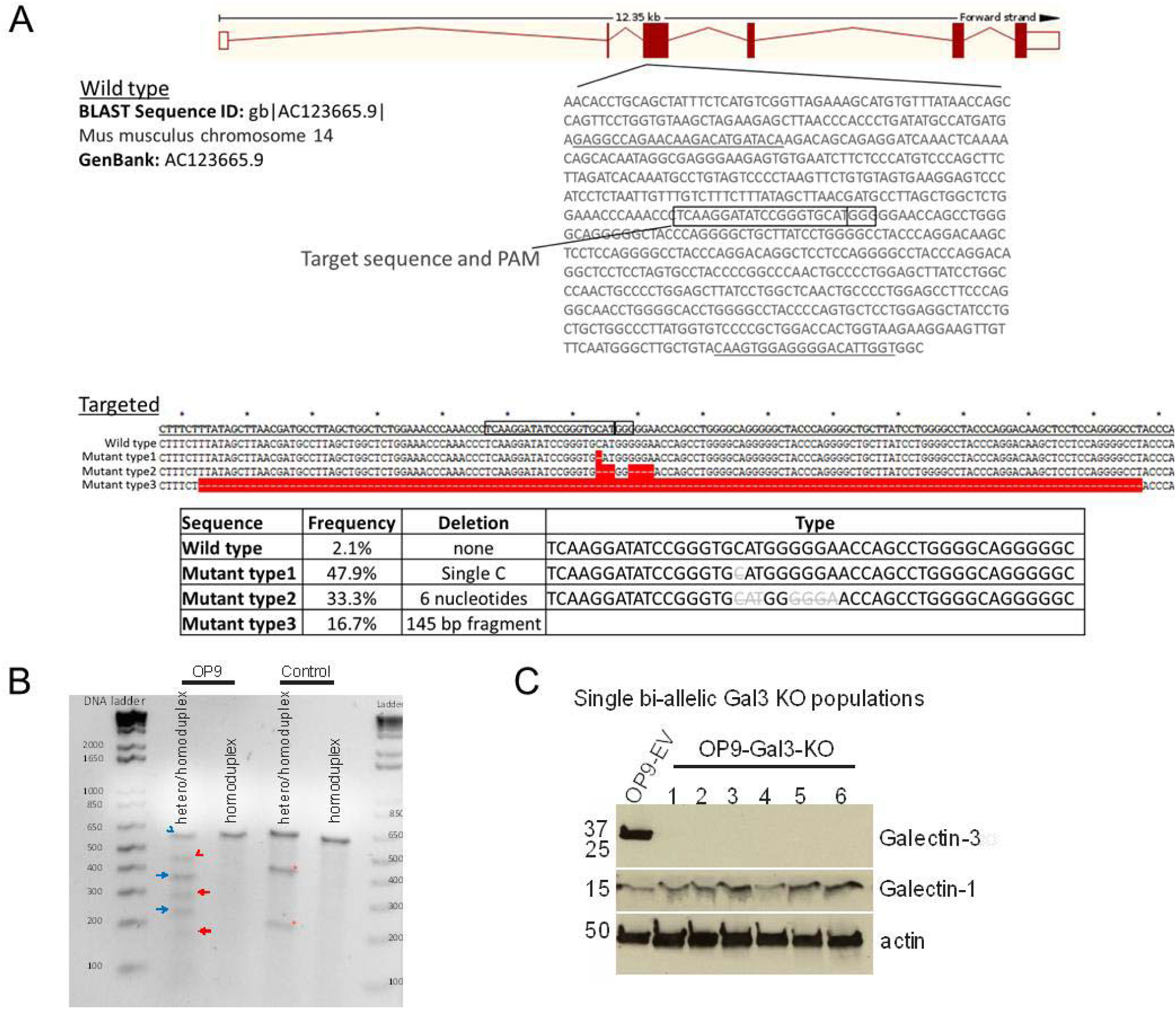
CRISPR/Cas9 mediated genome editing to knockout Gal3 in OP9 cells. (A) The CRISPR/Cas9 target region in exon 3 of the *lgals3* gene. The OP9-Gal3-KO has nucleotide deletion compared to the wild-type gene; this leads to a frameshift missense mutation and premature stop codons downstream in exon 3. (B) Surveyor assay to detect DNA breaks induced by CRISPR/Cas9 in exon 3 of OP9 cells. (C) Western blot showing Gal3 knockout in OP9 clones carrying bi-allelic CRISPR-induced mutation.

**Figure S3.**
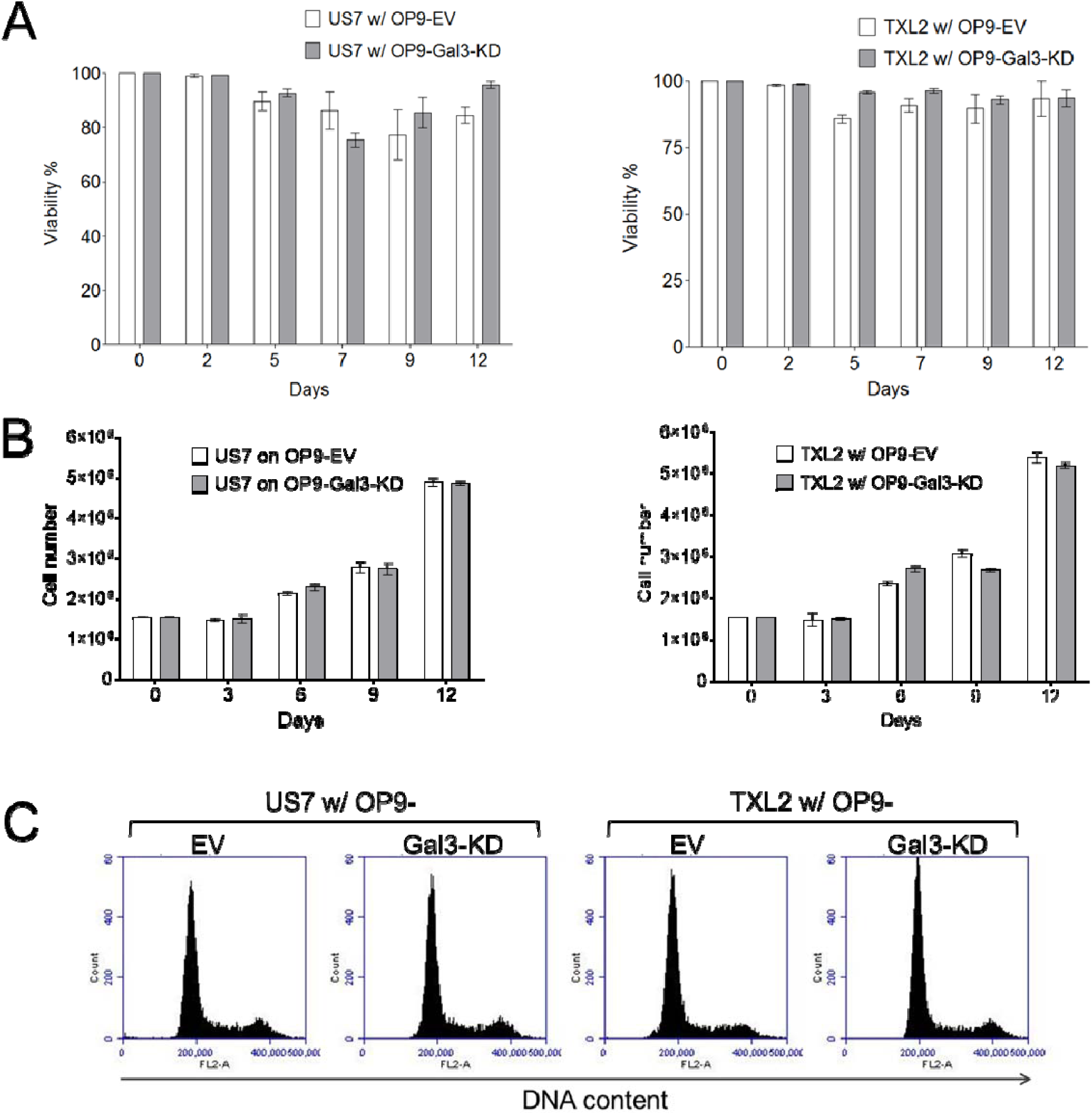
Under steady-state growth, Galectin-3 produced by stromal cells is not essential. (A) viability and (B) cell counts of US7 [left] and TXL2 [right] cells measured by Trypan blue exclusion grown over 12 days on the indicated OP9 stromal cells. (C) cell cycle analyzed by FACS and DNA content at different phases of the cell cycle in US7 or TXL2 cells grown for more than 4 days in co-culture with the indicated OP9 cells.

**Figure S4.**
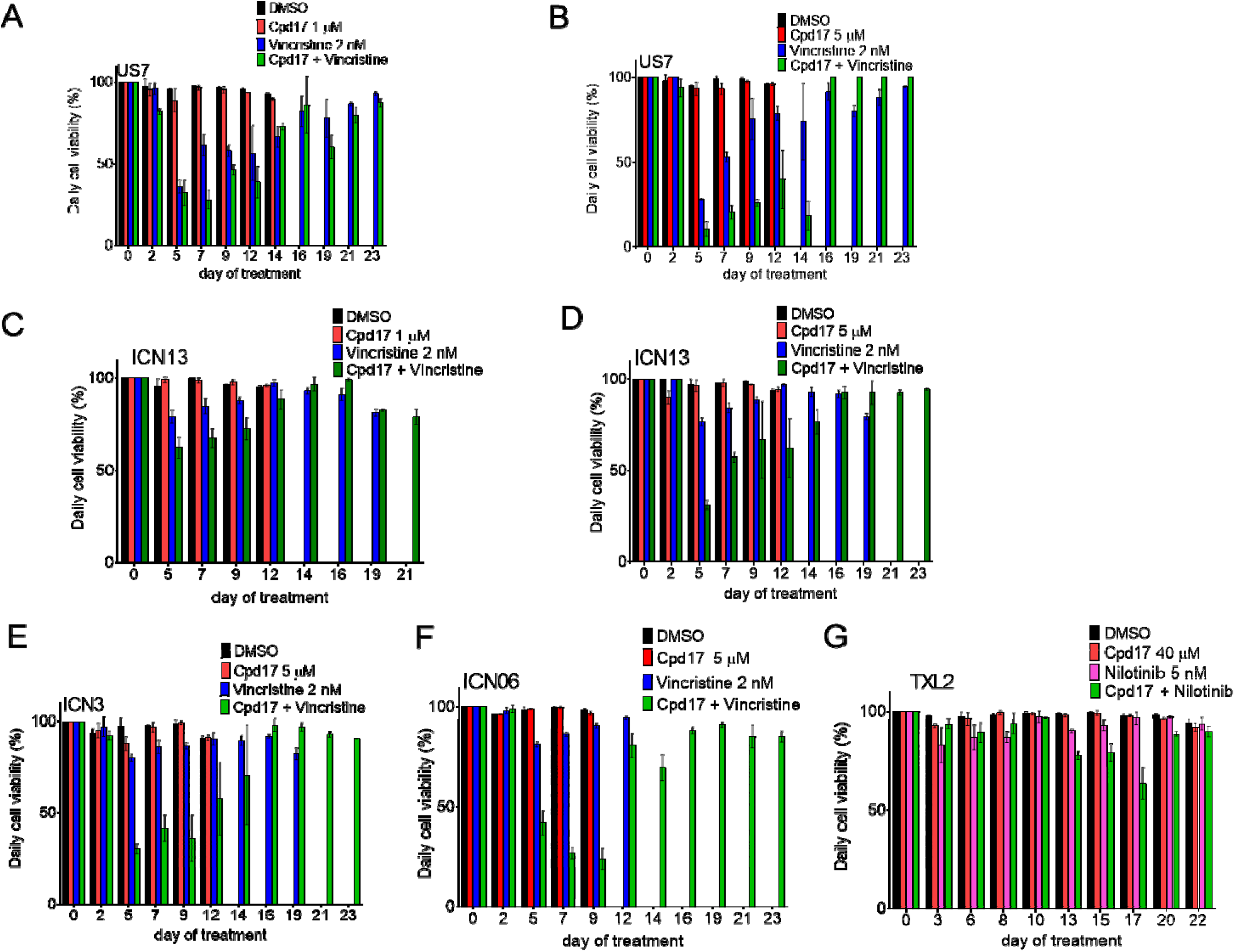
Viability of BCP-ALLs treated with combination treatment with Cpd17. Samples (Fig. 8) were treated with solvent DMSO [black bars], Cpd17 [red bars], standard chemotherapy [blue bars] or a combination of chemotherapy with Cpd17, [green bars]. DMSO only, Cpd17 mono-treatment and vincristine mono-treatment samples were not followed after a certain period when cell numbers exceeded the capacity of the wells and cultures became overgrown in A-F. Viability = viable cell count/total cell count x 100%. This is a measurement for the Trypan-blue excluding [living] status of cells that remain in the culture, even if they are only very few in number. Error bars, mean ±SEM of triplicate wells. Fresh drugs at the same concentration were added with every medium change. However, in panel G, Cpd17 was administered at 10 μM on day 0, increased to 20 μM on day 6 and further increased to 40 μM on day 10 because drug combination effects were minimal at 10 and 20 μM. A-F: 2 nM vincristine as standard chemotherapy; G: 5 nM targeted tyrosine kinase inhibitor nilotinib. A-B: US7 cells; C-D: ICN13; E: ICN3; F: ICN06; G: TXL2. A, C: 1 μM Cpd17. B, D-F: 5 μM Cpd17. G: 40 μM Cpd17 from d10 onwards.

